# A cup of coffee can boost your motor performance: Enhanced motor sequence learning with caffeine

**DOI:** 10.1101/2024.08.22.609116

**Authors:** Hakjoo Kim, David L. Wright, Jamie Kweon, Joshua C. Brown

## Abstract

Caffeine is consumed in various beverages, such as coffee, energy drinks, soda, and tea. The effects of caffeine have been shown to manifest in various ways across different studies. Some studies suggest that caffeine can enhance motor performance. Despite this, numerous motor learning studies do not control for participants’ caffeine intake or do not record whether participants consumed caffeine. The purpose of this study was to compare the motor learning performance (i.e., online improvement) of individuals who consumed caffeine and those who did not before practice with a novel motor sequence task. Sixty-five right-handed healthy undergraduate students participated in this study. Individuals reported their caffeine consumption through a pre-experiment questionnaire prior to performing an eight-item serial reaction time task. The participants who consumed caffeine reported drinking one cup of coffee on average 158.8 minutes before performing the sequential task. The motor performance of individuals who consumed caffeine was compared to those who did not, with the caffeine-consuming group (*n* = 13) exhibiting faster response times during online learning than the non-caffeine group (*n* = 52). The findings of this study suggest that caffeine should be controlled for in future motor learning research.

## Introduction

Caffeine (1,3,7-trimethylpurine-2,6-dione) is the most widely used stimulant of the central nervous system, belonging to the methylxanthine group (Nehlig et al., 1992). A variety of commonly consumed beverages, including coffee, tea, energy drinks, and soft drinks, contain caffeine (McLellan et al., 2016; Roberts, 2021). Caffeine is rapidly absorbed following ingestion, and although there is individual variation, it typically takes about 30 to 90 minutes to reach peak plasma concentration (Newton et al., 1981). The half-life of caffeine in plasma can differ significantly between individuals, as Blanchard and Sawers (1983) reported a range of 2.7 to 9.9 hours.

Caffeine is a competitive (Phillis et al., 1979) and non-selective adenosine receptor antagonist (Fredholm et al., 1999; McLellan et al., 2016). Caffeine has a similar chemical structure to adenosine, allowing it to bind to and block adenosine receptors, mainly the A_1_ and A_2A_ receptors. When adenosine binds to its receptors, it exerts inhibitory effects, leading to fatigue (Lorist & Tops, 2003) and drowsiness (Porkka-Heiskanen et al., 1997). However, when caffeine binds to adenosine receptors, particularly A_2A_ receptors, it blocks these inhibitory effects, thereby increasing alertness and inducing arousal (Huang et al., 2005; Lazarus et al., 2011). In addition, when adenosine binds to A_2A_ receptors, it inhibits dopamine signaling. Therefore, when caffeine competitively binds to A_2A_ receptors, it blocks adenosine’s inhibitory effects and results in increased dopamine release (Ferré et al., 1992). Caffeine is also an antagonist of benzodiazepine receptors, part of the γ-Aminobutyric acid type A (GABA_A_) receptor complex. However, it has a weak binding affinity compared to adenosine receptors, inhibiting GABA_A_ receptors only at toxic levels of caffeine (Fredholm et al., 1999; McLellan et al., 2016; Nehlig et al., 1992; Reddy et al., 2024; Song et al., 2023).

Caffeine may affect synaptic plasticity, potentially influencing the strength and adaptability of connections between neurons in the brain. It has been observed that long-term potentiation (LTP), regarded as a key mechanism underlying learning and memory formation (Morris, 1989; Bliss & Collingridge, 1993; Nabavi et al., 2014; Brown et al., 2022), is promoted by selective adenosine A_1_ receptor antagonists, while selective adenosine A_2A_ receptor antagonists attenuate LTP (Lu et al., 1999; de Mendoça & Ribeiro, 2001; Costenla et al., 2010). Several studies have explored whether caffeine affects synaptic plasticity using non-invasive brain stimulation techniques, such as transcranial magnetic stimulation (TMS), in humans (Hanajima et al., 2019; Orth et al., 2005; Vigne et al., 2023; Zulkifly et al., 2020). Orth et al. (2005) reported that 3 mg/kg body weight of caffeine did not influence cortical excitability (i.e., resting and active motor thresholds, intra-cortical facilitation, short-interval intracortical inhibition, and cortical silent periods). However, Hanajima et al. (2019) revealed that consuming 200 mg of caffeine reduces LTP-like plasticity induced by quadripulse TMS. Similarly, Vigne et al. (2023) suggested that chronic caffeine users may have a reduced capacity for LTP-like plasticity, induced by 10-Hz repetitive TMS, compared to non-caffeine users. The results of Zulkifly et al. (2020) using transcranial alternating current stimulation also demonstrated that drinking espresso decreases LTP-like plasticity.

Beyond its influence on synaptic plasticity, numerous studies have shown that caffeine consumption can enhance motor performance (see McLellan et al., 2016; Pesta et al., 2013). For example, an early study reported that small doses of caffeine (i.e., 1-3 grains) accelerated typewriting speed, while larger doses (i.e., 4-6 grains) reduced performance (Hollingworth, 1912). Decades later, LS Miller and SE Miller (1996) investigated the effects of different amounts of caffeine on motor performance of a multiple-force discrimination task. In their study, participants took a placebo, 1 mg, 3 mg, or 5 mg of caffeine per kg body weight approximately 45 minutes prior to the experiment. LS Miller and SE Miller revealed that individuals with 3 or 5 mg/kg showed faster responses during the initial sessions compared to those with placebo or 1 mg/kg. However, caffeine does not always have a positive effect on motor learning. For instance, Mednick et al. (2008) revealed that 200 mg of caffeine administered 60 minutes prior to a five-item keyboard sequence task (Walker et al., 2002) significantly impaired the offline performance of motor sequence learning—referring to the retention or change in skills after practice. More recently, Hussain and Cole (2015) reported that 200 mg of caffeine administered after skill acquisition did not influence both the online learning—referring to real-time improvements during practice—and the offline learning of a continuous isometric visuomotor tracking task. However, research examining how caffeine consumption affects online performance of motor sequence learning is still limited.

The purpose of the present study is to investigate the effect of caffeine consumption on motor sequence learning. Based on numerous studies that have demonstrated the positive effects of caffeine on motor performance, our hypothesis is that caffeine consumption would enhance the online performance of motor sequence learning. Should this study demonstrate that caffeine influences motor learning, it could serve as evidence that caffeine intake needs to be controlled in future motor learning research and could warrant further investigation into optimization of motor learning and performance.

## Results

### Overall performance

The 2 (Group: No Caffeine, Caffeine) x 15 (Trial: 1 to 15) repeated measures analysis of variance (ANOVA) performed on the mean response time (RT) data revealed a significant main effect of Trial, *F*(7.898, 497.547) = 37.358 (Greenhouse-Geisser), *p* < .001, partial *η*^2^ = .372 (see Figure 1). However, there were no effects of Group, *F*(1, 63) = 0.535, *p* = .467, partial *η*^2^ = .008, and the interaction between Group and Trial, *F*(7.898, 497.547) = 1.676 (Greenhouse-Geisser), *p* = .103, partial *η*^2^ = .026. In other words, as the amount of practice in the training sequence increased, participants’ performance improved (i.e., faster performance) regardless of caffeine condition.

**Figure 1.**
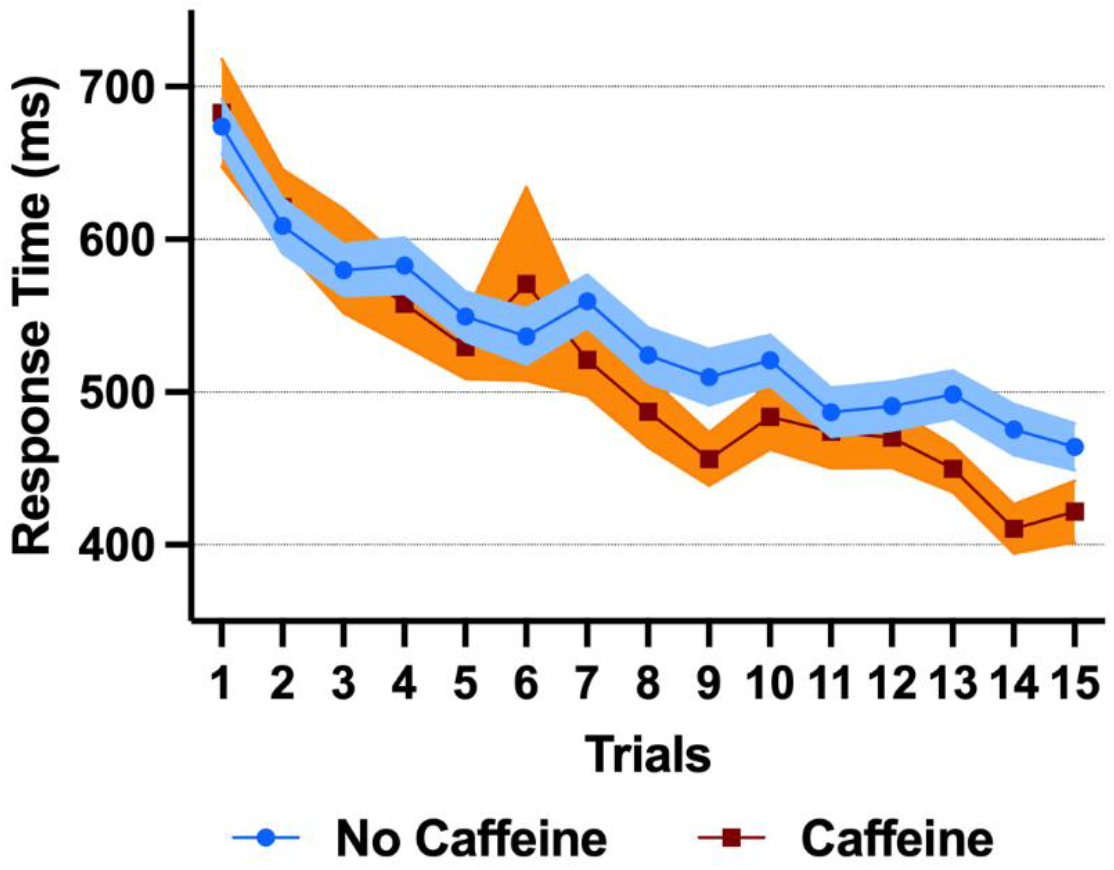
Overall task performance of the training sequence. The blue circles denote the No Caffeine group, and the brown squares denote the Caffeine group. The shaded area represents the standard errors.

### Online performance

The mean RT of the first three trials (i.e., Trials 1, 2, and 3) and the mean RT of the last three trials (i.e., Trials 13, 14, and 15) were utilized to analyze online performance improvement (Kim et al., 2024). RT ratio was calculated by dividing the mean RT of the last three trials by the mean RT of the first three trials, and the ratio data were submitted to an independent samples *t*-test (Group: No Caffeine, Caffeine) to examine the effect of caffeine consumption on online gain. Levene’s test for equality of variances confirmed that the group variances were equal (*p* = .406). The independent samples *t*-test revealed that there was a difference between the No Caffeine (M = 0.774, SEM = 0.014) and Caffeine (M = 0.685, SEM = 0.026) groups, *t*(63) = 2.907, *p* = .005 (see Figure 2). This indicates that the individuals who consumed caffeine demonstrated greater improvement in response time compared to those who did not.

**Figure 2.**
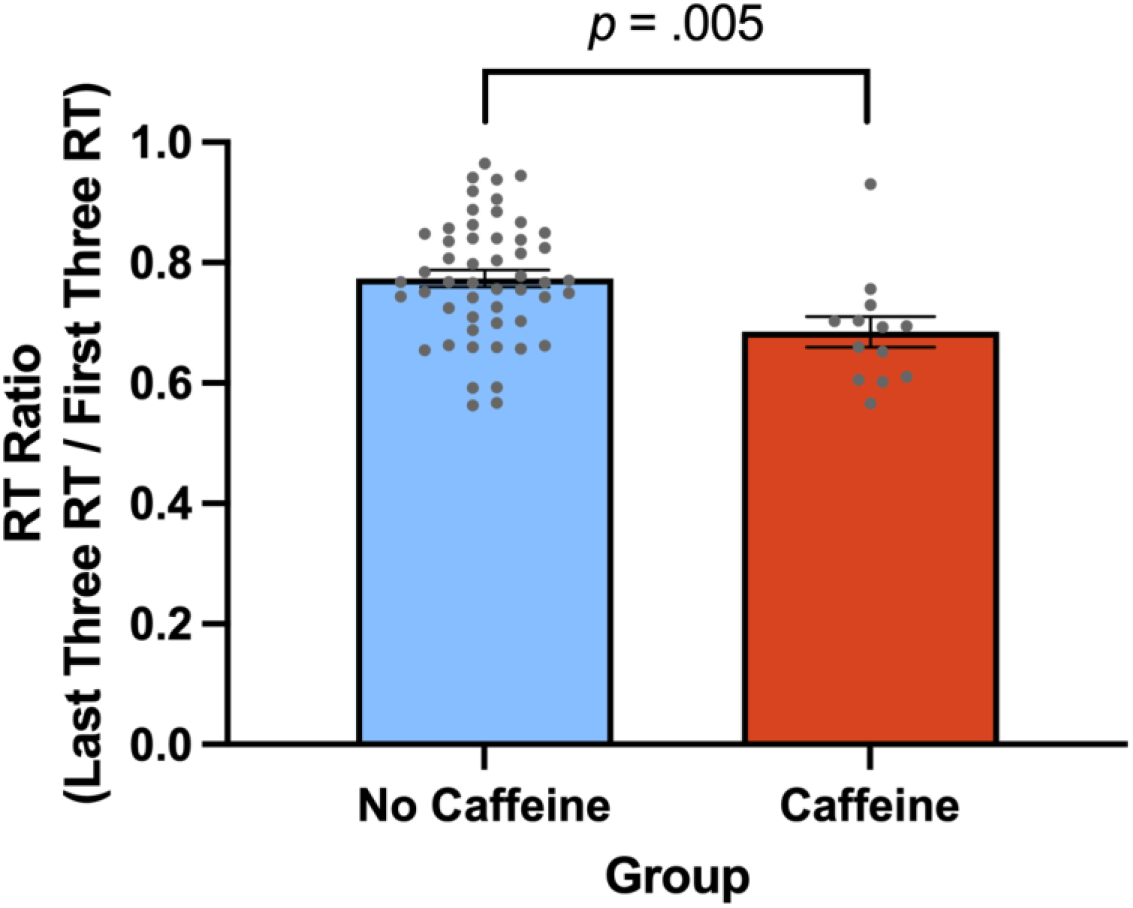
Response time (RT) ratio in the No Caffeine and Caffeine groups. The RT ratio was calculated by taking the mean RT of the last three trials (i.e., Trials 13, 14, and 15) and dividing it by the mean RT of the first three trials (i.e., Trials 1, 2, and 3). The gray points represent the individual data in the No Caffeine (*n* = 52) and Caffeine groups (*n* = 13). The smaller the RT ratio, the greater the performance improvement. The error bars represent the standard errors.

**Figure 3.**
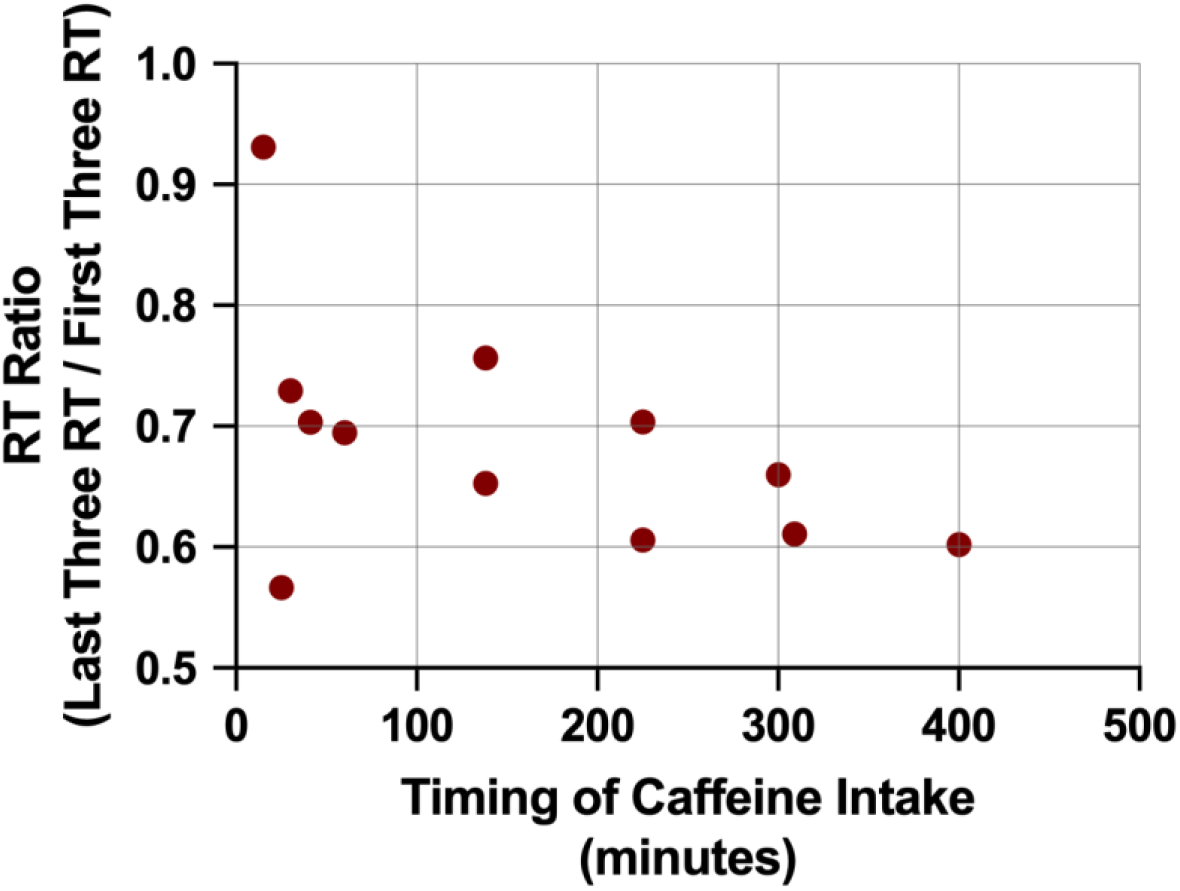
Correlation between online performance and timing of caffeine intake. The smaller the RT ratio, the greater the performance improvement. The X-axis represents when the participant consumed caffeine prior to the experiment. For example, ‘60 (minutes)’ indicates that caffeine was consumed 60 minutes before participating in the experiment.

### Online performance and timing of caffeine consumption

Among the 65 participants, 20% (i.e., 13 participants) reported that they consumed caffeine from a cup (i.e., ≥ 12 ounces) of caffeinated Americano coffee before participating in the experiment. Of these 13 participants, 12 individuals were asked when they had drunk the coffee, and average coffee consumption was 158.83 minutes (SD: 131.12 minutes) prior to the experiment. The relationship between the timing of coffee intake and online performance gain was analyzed using Pearson’s correlation coefficient. The analysis indicated a moderate negative correlation (*r* = -.487). However, the correlation was not statistically significant (*p* = .109). Thus, the current data did not provide a clear indication of how the timing of caffeine consumption affects online improvement.

## Discussion

The current study revealed that caffeine consumption (i.e., a cup of coffee) facilitated the online improvement of motor sequence learning. These findings are consistent with previous studies that demonstrated reductions in RT and benefits to motor performance from caffeine intake (for reviews, see McLellan et al., 2016; Pesta et al., 2013). However, the mechanisms underlying the effects of caffeine on motor learning remain unclear. While this study found an effect of caffeine on motor learning, the studies by Mednick et al. (2008) and Hussain and Cole (2015) did not find an effect of caffeine on improving task performance. One of the differences between the present study and those studies was the nature of the feedback the participants were provided during training. Mednick et al. used a white dot that appeared on the screen to indicate the response to be made instead of a number, which was associated with a particular key press, to avoid providing feedback on accuracy (also see Walker et al., 2002). In contrast, Hussain and Cole provided continuous visual feedback regarding the cursor position, which is assumed to be central to the use of an error-based learning strategy (see Krakauer et al., 2019; Spampinato & Celnik, 2021). However, in the present study, the feedback provided to participants was more in keeping with that commonly used to encourage reinforcement learning, a learning strategy considered to be distinct from error-based learning. Similarly, in the study by LS Miller and SE Miller (1996), individuals received auditory and visual feedback regarding the accuracy of their responses, which again was more akin to reinforcement learning. Reinforcement learning is a process in which learning occurs by rewarding correct responses (i.e., key press), which has been associated with modulating dopaminergic activity (Krakauer et al., 2019; Spampinato & Celnik, 2021). The increased dopamine levels due to caffeine consumption (Ferré et al., 1992) may have facilitated reinforcement learning. However, it cannot be ruled out that the motor performance improvement was simply due to the arousal effect of caffeine, allowing participants to focus better on the motor sequence task. Nonetheless, in the study by Lin et al. (2023), no difference in response time was observed between the caffeine and placebo groups in a simple recognition-response task (i.e., 0-back task) that tested attention and motor control without challenging the memory capacity. Future research should compare the effects of caffeine across different learning strategies.

Despite the outcomes, this study has several limitations. First, the caffeine data were collected through participants’ self-reported surveys based on their memory. Therefore, there is a possibility that individuals may have incorrectly reported not consuming coffee, perhaps forgetting that they had actually consumed it (or vice versa). However, the accuracy of the participants’ responses could not be verified in this study. Second, although individuals in the Caffeine group reported consuming a cup (i.e., ≥ 12 ounces) of caffeinated Americano coffee, the exact amount of caffeine contained in that single cup was not measured. Third, while the timing of caffeine intake before the experiment was recorded, it varied among participants. Although the correlation analysis of our data was not statistically significant, considering that peak plasma concentration is generally reached approximately 30-90 minutes after consumption (Newton et al., 1981), it is possible that the effects of caffeine could vary significantly depending on the timing of its intake. Additionally, while we collected the time individuals started drinking coffee, we do not know the time they finished. Future studies should administer a consistent dose of caffeine at a fixed time to compare its effects. Lastly, while this study compared the effects of caffeine on the online improvement of motor sequence learning, future research should investigate a broader examination of the influence of caffeine on learning, focusing on potential impacts on offline improvement and memory consolidation. Despite the limitations mentioned above, this study suggests that caffeine should be considered a potential controlled factor in motor learning research.

## Methods

### Participants

Sixty-five right-handed healthy undergraduate students (mean age ± SD: 20.23 ± 1.52, age range: 18-24, 43 females and 22 males) from the Department of Kinesiology and Sport Management at Texas A&M University participated in the current study. Written informed consent approved by the Texas A&M University Institutional Review Board was obtained from all participants. None of the participants were professional musicians. All participants received extra credit for an undergraduate Kinesiology course after completing the experiment. Table 1 shows the demographic information of the participants in each group.

**Table 1.** Demographic information. kg: kilograms, SD: standard deviation.

| Group | *n* | Age (SD) | Weight in kg (SD) | Females | Males |
| --- | --- | --- | --- | --- | --- |
| No Caffeine | 52 | 20.15 (1.47) | 69.63 (14.16) | 32 | 20 |
| Caffeine | 13 | 20.54 (1.71) | 72.75 (12.41) | 11 | 2 |
| Total | 65 | 20.23 (1.52) | 70.25 (13.79) | 43 | 22 |

### Caffeine intake

In this study, caffeine was not directly administered to participants at the laboratory. Instead, information about the type, timing, and amount of caffeine they consumed prior to the experiment was collected through a pre-experiment questionnaire. However, a survey on participants’ chronic caffeine use and dependence was not conducted. Individuals’ salivary, plasma, and urinary caffeine levels were not obtained.

### Serial reaction time task (SRTT)

An eight-item serial reaction time task (SRTT) was utilized to measure motor sequence learning. The SRTT was identical to the one used in Kim et al. (2024). In this study, however, only a single practice session was performed. Participants placed the four fingers (i.e., pinky, ring, middle, and index fingers) of their left hand on the V, B, N, and M keys of a standard computer keyboard. The V, B, N, and M keys corresponded to the numbers 1, 2, 3, and 4, respectively. Individuals repeatedly entered a specific sequence (i.e., 4-3-1-2-4-2-1-3) explicitly displayed in a row on the computer monitor for 30 seconds. In addition to the training sequence, there were five random sequences (i.e., Random #1: 4-1-3-2-3-1-2-4, Random #2: 2-4-3-1-3-1-4-2, Random #3: 3-4-2-1-3-2-4-1, Random #4: 1-4-3-2-1-2-4-3, and Random #5: 4-2-3-1-2-4-1-3), which appeared in the first trial and following every three trials of the training sequence (i.e., R-T-T-T-R-T-T-T-R-T-T-T-R-T-T-T-R-T-T-T; R: random sequence, T: training sequence; see Kim et al., 2024). After each 30-second trial, a 15-second rest period was provided. During the 15-second rest period, participants could not see any sequences on the monitor. While typing the training or random sequences, individuals received real-time feedback, such as the number of sequences they correctly entered in each trial and whether they pressed the keys accurately. Participants were instructed to increase this number as much as possible. The SRTT was performed for a total of 15 minutes (= 20 trials × (30 seconds/trial + 15 seconds/rest) = 900 seconds).

### Procedure

After submitting informed consent and pre-experiment questionnaires, participants sat at the computer and received instructions about the SRTT from the experimenter. They then performed only one session of the SRTT using their left hand for 15 minutes. All experimental procedures were completed within one hour.

### Data analysis

The dependent variable of the current study was the mean RT (in milliseconds) for a correct key press of the training sequence. In this study, the random sequences’ data were not analyzed. Statistical analyses, such as two-way ANOVA and independent samples *t*-test, were conducted using SPSS software (Version 28.0.0.0, IBM, United States). Normality was assessed using the Shapiro-Wilk test, and outliers were not removed.

## Author contributions

HK: Conceptualization Data curation, Formal analysis, Funding acquisition, Methodology, Validation, Visualization, Writing – original draft, Writing – review & editing; JK: Formal analysis, Validation, Writing – original draft, Writing – review & editing; JCB: Formal analysis, Validation, Writing – original draft, Writing – review & editing; DLW: Conceptualization, Data curation, Formal analysis, Methodology, Project administration, Resources, Supervision, Validation, Writing – original draft, Writing – review & editing.

## Acknowledgements

This work was supported by the School of Education & Human Development (SEHD) Research Grant at Texas A&M University that was awarded to HK.

## Data availability

The datasets analyzed during the current study are available from the corresponding author on reasonable request.

## Competing interests

The authors declare no competing interests.

## Notes

### Competing Interest Statement

The authors have declared no competing interest.

## References

Blanchard, J., & Sawers, S. J. A. (1983). The absolute bioavailability of caffeine in man. European Journal of Clinical Pharmacology, 24, 93–98. 10.1007/BF00613933

Bliss, T. V., & Collingridge, G. L. (1993). A synaptic model of memory: long-term potentiation in the hippocampus. Nature, 361(6407), 31–39. 10.1038/361031a0

Brown, J. C., Higgins, E. S., & George, M. S. (2022). Synaptic plasticity 101: the story of the AMPA receptor for the brain stimulation practitioner. Neuromodulation, 25(8), 1289–1298. 10.1016/j.neurom.2021.09.003

Costenla, A. R., Cunha, R. A., & De Mendonça, A. (2010). Caffeine, adenosine receptors, and synaptic plasticity. Journal of Alzheimer’s Disease, 20(S1), S25–S34. 10.3233/jad-2010-091384

de Mendonça, A., & Ribeiro, J. A. (2001). Adenosine and synaptic plasticity. Drug Development Research, 52(1-2), 283–290. 10.1002/ddr.1125

Ferré, S., Fuxe, K., Von Euler, G., Johansson, B., & Fredholm, B. B. (1992). Adenosine-dopamine interactions in the brain. Neuroscience, 51(3), 501–512. 10.1016/0306-4522(92)90291-9

Fredholm, B. B., Bättig, K., Holmén, J., Nehlig, A., & Zvartau, E. E. (1999). Actions of caffeine in the brain with special reference to factors that contribute to its widespread use. Pharmacological Reviews, 51(1), 83–133.

Hanajima, R., Tanaka, N., Tsutsumi, R., Shirota, Y., Shimizu, T., Terao, Y., & Ugawa, Y. (2019). Effect of caffeine on long-term potentiation-like effects induced by quadripulse transcranial magnetic stimulation. Experimental Brain Research, 237, 647–651. 10.1007/s00221-018-5450-9

Hollingworth, H. L. (1912). The influence of caffeine on the speed and quality of performance in typewriting. Psychological Review, 19(1), 66. https://psycnet.apa.org/doi/10.1037/h0073935

Huang, Z. L., Qu, W. M., Eguchi, N., Chen, J. F., Schwarzschild, M. A., Fredholm, B. B., Urade, Y., & Hayaishi, O. (2005). Adenosine A2A, but not A1, receptors mediate the arousal effect of caffeine. Nature Neuroscience, 8(7), 858–859. 10.1038/nn1491

Hussain, S. J., & Cole, K. J. (2015). No enhancement of 24-hour visuomotor skill retention by post-practice caffeine administration. PLoS ONE, 10(6), e0129543. 10.1371/journal.pone.0129543

Kim, H., King, B., Verwey, W. B., Buchanan, J. J., & Wright, D. L. (2024). Timing of transcranial direct current stimulation at M1 does not affect motor sequence learning. Heliyon, 10(4), E25905. 10.1016/j.heliyon.2024.e25905

Krakauer, J. W., Hadjiosif, A. M., Xu, J., Wong, A. L., & Haith, A. M. (2019). Motor learning. Comprehensive Physiology, 9(2), 613–663. 10.1002/cphy.c170043

Lazarus, M., Shen, H. Y., Cherasse, Y., Qu, W. M., Huang, Z. L., Bass, C. E., … & Chen, J. F. (2011). Arousal effect of caffeine depends on adenosine A2A receptors in the shell of the nucleus accumbens. Journal of Neuroscience, 31(27), 10067–10075. 10.1523/JNEUROSCI.6730-10.2011

Lin, Y. S., Weibel, J., Landolt, H. P., Santini, F., Slawik, H., Borgwardt, S., … & Reichert, C. F. (2023). Brain activity during a working memory task after daily caffeine intake and caffeine withdrawal: a randomized double-blind placebo-controlled trial. Scientific Reports, 13(1), 1002. 10.1038/s41598-022-26808-5

Lorist, M. M., & Tops, M. (2003). Caffeine, fatigue, and cognition. Brain and Cognition, 53(1), 82–94. 10.1016/S0278-2626(03)00206-9

Lu, K. T., Wu, S. P., & Gean, P. W. (1999). Promotion of forskolin-induced long-term potentiation of synaptic transmission by caffeine in area CA1 of the rat hippocampus. Chinese Journal of Physiology, 42(4), 249–253.

McLellan, T. M., Caldwell, J. A., & Lieberman, H. R. (2016). A review of caffeine’s effects on cognitive, physical and occupational performance. Neuroscience & Biobehavioral Reviews, 71, 294–312. 10.1016/j.neubiorev.2016.09.001

Mednick, S. C., Cai, D. J., Kanady, J., & Drummond, S. P. (2008). Comparing the benefits of caffeine, naps and placebo on verbal, motor and perceptual memory. Behavioural Brain Research, 193(1), 79–86. 10.1016/j.bbr.2008.04.028

Miller, L. S., & Miller, S. E. (1996). Caffeine enhances initial but not extended learning of a proprioceptive-based discrimination task in nonsmoking moderate users. Perceptual and Motor Skills, 82(3), 891–898. 10.2466/pms.1996.82.3.891

Morris, R. G. (1989). Synaptic plasticity and learning: selective impairment of learning rats and blockade of long-term potentiation in vivo by the N-methyl-D-aspartate receptor antagonist AP5. Journal of Neuroscience, 9(9), 3040–3057. 10.1523/JNEUROSCI.09-09-03040.1989

Nabavi, S., Fox, R., Proulx, C. D., Lin, J. Y., Tsien, R. Y., & Malinow, R. (2014). Engineering a memory with LTD and LTP. Nature, 511(7509), 348–352. 10.1038/nature13294

Nehlig, A., Daval, J. L., & Debry, G. (1992). Caffeine and the central nervous system: mechanisms of action, biochemical, metabolic and psychostimulant effects. Brain Research Reviews, 17(2), 139–170. 10.1016/0165-0173(92)90012-B

Newton, R. L. M. H. I., Broughton, L. J., Lind, M. J., Morrison, P. J., Rogers, H. J., & Bradbrook, I. D. (1981). Plasma and salivary pharmacokinetics of caffeine in man. European Journal of Clinical Pharmacology, 21, 45–52. 10.1007/BF00609587

Pesta, D. H., Angadi, S. S., Burtscher, M., & Roberts, C. K. (2013). The effects of caffeine, nicotine, ethanol, and tetrahydrocannabinol on exercise performance. Nutrition & Metabolism, 10, 1–15. 10.1186/1743-7075-10-71

Phillis, J. W., Edstrom, J. P., Kostopoulos, G. K., & Kirkpatrick, J. R. (1979). Effects of adenosine and adenine nucleotides on synaptic transmission in the cerebral cortex. Canadian Journal of Physiology and Pharmacology, 57(11), 1289–1312. 10.1139/y79-194

Porkka-Heiskanen, T., Strecker, R. E., Thakkar, M., Bjørkum, A. A., Greene, R. W., & McCarley, R. W. (1997). Adenosine: a mediator of the sleep-inducing effects of prolonged wakefulness. Science, 276(5316), 1265–1268. 10.1126/science.276.5316.1265

Roberts, A. (2021). Caffeine: an evaluation of the safety database. In R. C. Gupta, R. Lall, & A. Srivastava (Eds.), Nutraceuticals (2nd ed., pp. 501–518). Academic Press. 10.1016/B978-0-12-821038-3.00032-X

Song, X., Kirtipal, N., Lee, S., Malý, P., & Bharadwaj, S. (2023). Current therapeutic targets and multifaceted physiological impacts of caffeine. Phytotherapy Research, 37(12), 5558–5598. 10.1002/ptr.8000

Spampinato, D., & Celnik, P. (2021). Multiple motor learning processes in humans: defining their neurophysiological bases. The Neuroscientist, 27(3), 246–267. 10.1177/1073858420939552

Vigne, M., Kweon, J., Sharma, P., Greenberg, B. D., Carpenter, L. L., & Brown, J. C. (2023). Chronic caffeine consumption curbs rTMS-induced plasticity. Frontiers in Psychiatry, 14, 1137681. 10.3389/fpsyt.2023.1137681

Walker, M. P., Brakefield, T., Morgan, A., Hobson, J. A., & Stickgold, R. (2002). Practice with sleep makes perfect: sleep-dependent motor skill learning. Neuron, 35(1), 205–211. 10.1016/S0896-6273(02)00746-8

Zulkifly, M. F. M., Merkohitaj, O., & Paulus, W. (2020). Transcranial alternating current stimulation induced excitatory aftereffects are abolished by decaffeinated espresso and reversed into inhibition by espresso with caffeine. Clinical Neurophysiology, 131(3), 778–779. 10.1016/j.clinph.2019.11.062

